# *Ardisia tchimbele sp. nov*. (Primulaceae) - A Critically Endangered forest shrub, and notes on the Crystal Mountains of Gabon

**DOI:** 10.64898/2026.11.19.767889

**Authors:** Harry Murdoch, Martin Cheek

**Affiliations:** Herbarium, Royal Botanic Gardens, Kew, Richmond, Surrey, TW9 3AE, UK

**Keywords:** Conservation, extinction, hydroelectric projects, oil glands

## Abstract

*Ardisia tchimbele sp. nov*. (Primulaceae) an erect, branching shrub, is described from the forests downstream of Tchimbélé, a hydroelectric installation in the Crystal Mountains, Gabon that supplies Libreville, the capital with energy. The new species is based on three specimens collected in the 1980s and 1990s. It keys out and has similarities with a taxon informally described in 1979 by Taton as *Ardisia* sp. 1, also from the Crystal Mts and which is slightly discordant morphologically.

The new species is considered threatened by the risk of flooding of its habitat from further hydroelectric projects and is provisionally assessed as Critically Endangered using the 2012 IUCN standard.

We offer as context a short review of the vegetation, topography and plant diversity of the Crystal Mountains of Gabon.

## Introduction

The genus *Ardisia* Sw. (Primulaceae, formerly Myrsinaceae) is pantropical, with 741 species accepted (POWO continuously updated). In continental Africa, the c. 50 described species are most diverse in Cameroon and Gabon and absent from West, East and South-Central Africa apart from a few species in western Uganda, eastern Nigeria and another in northern metropolitan Angola (Peng & Cheek 2025). An introduction to the taxonomy, conservation, chemistry and uses of the genus, and the discovery of species in continental Africa can be found in Peng & Cheek (2025). Additional uses of Cameroon *Ardisia* species can be found in Pollard et al. (2026).

This paper is one of several preparatory to the Flore Du Gabon account of *Ardisia*. Following a revision (Peng & Cheek 2025) of both Gabonese and Cameroonian species of the herbaceous chamaephyte group (see African *Ardisia* life form classification, Cheek *et al*. 2025) and of the Gabonese monocaul phanerophyte species (Murdoch & Cheek 2026a), together with an additional Gabonese herbaceous species (Murdoch & Cheek 2026b), we here describe a new species in the branched phanerophyte (shrubby) group, the most species-diverse of the four *Ardisia* groups in tropical Africa (Cheek *et al*. 2025). Eleven species of this group are currently accepted for Gabon (Sosef et al. 2006) which were last revised in the monumental account of Taton (1979). Since Taton (1979) no additional new species of this group have been described from Gabon, although two species have been described from the mountains of western Cameroon (Cheek in Cheek 2010).

The three specimens of the proposed new species were found to match no other species known from Gabon, nor elsewhere in tropical Africa. The materials, including both flowers and fruits, were all collected from 1989 to 1994, from near Tchimbélé in the Crystal Mts of Gabon. The specimens can be distinguished from those of all other shrubby species of the genus in tropical Africa by the comparatively large size of the oblong-elliptic, entire leaf blades (length to breadth ratio 2.6-3.4: 1) which are further remarkable for the high density and highly conspicuous bright orange-red colour (transmitted light and hand lense needed) of the internal oil glands, coupled with the high density of red scales (visible with x 10 hand lense) on the abaxial blade surface. In the key to *Ardisia* of tropical Africa (Taton 1979) this taxon keys out as *Ardisia* sp. 1 (see NOTES below) due to the characters mentioned above and the axillary, pseudo-fasciculate inflorescences in which the bracts are scale-like, the sepals ovate-acute. We here formally describe this taxon, compare it with the species which are most similar, and assess its risk of extinction.

## Materials and Methods

Specimens and measurements were prepared/examined following the protocols outlined in Peng & Cheek (2025) and M. All specimens cited have been seen.

### Taxonomic Treatment

#### Ardisia tchimbele

Cheek & Murdoch **sp. nov**. Fig. 1

**Fig. 1.**
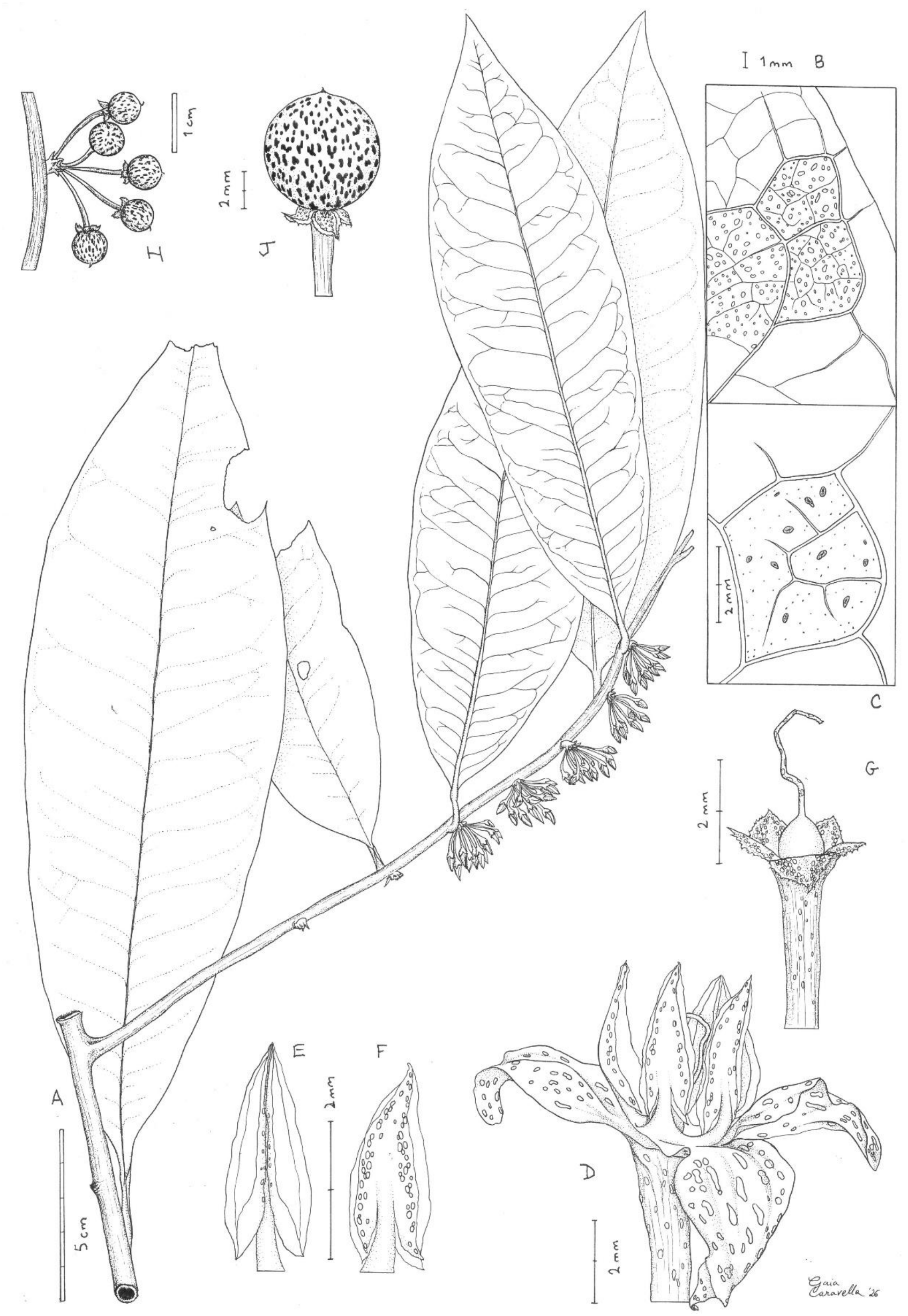
*Ardisia tchimbele* **A** habit, orthotropic and flowering plagiotropic stems; **B** adaxial leaf blade surface near margin showing the areolae with oil glands (unshaded); **C** an areolae of abaxial blade surface showing 11 elliptic oil glands (shown dotted) and numerous smaller surface scales (shown in black) **D** flower at anthesis, side view, hydrated; **E** stamens, ventral, adaxial (left) and abaxial, dorsal views (right); **F** flower showing gynoecium and calyx (corolla removed); **G** infructescence (note peduncle); **J** fruit, sepals and distal pedicel. **A-G** from *Breteler* 12910 (WAG, holo.), **H-J** from *Wieringa* 248 (WAG). All drawn by Gaia Caravella.

- Type: Gabon, Woleu-Ntem, Crystal Mountains, near Tchimbélé. Old secondary forest, 0°36’N, 10°23’E, 650 m, b., fl., 17 Sep. 1994, *Breteler* 12910 with Nzabi & Wieringa (holotype WAG0040953; isotypes BR0000017904957; K006391366, MO6680737; WAG0040952).

Syn. *Ardisia mildbraedii* sensu Sosef *et al*. (2006) pro parte quoad. *J*.*J. Wieringa* 248! non (Gilg & Schellenb.) Taton (1979)

*Branching erect shrub, 0*.*5-1*.*1 m tall*. Main axis (orthotropic stem) vertical, phyllotaxy helical (spiral insertion), subterete, 5-6 mm diam., internodes c. 16 mm long, glabrescent, epidermis yellow brown to grey brown, “stem exuding orange juice” (*J*.*J. Wieringa* 248). Plagiotropic stems ±horizontal, phyllotaxy distichous, terete, stems 15-22 cm long, 2.5-3.5 mm diam., internodes 7-9, 14-70 mm long with longest internode proximal, often comprising half the length of the stem; stem at the attachment point to the main axis abruptly dilated 7-11 mm wide, and laterally flattened, indumentum and epidermis as orthotropic stem.

*Leaves* of orthotropic stem, petiole canaliculate, 9-22 × 2.5-3.5 mm; lamina discolorous, adaxially dark brown green, abaxially paler brown, coriaceous in life (*F*.*J. Breteler* 12910), elliptic-oblong, 19.1-28.6 × 7.4-9.2 cm, length to width ratio 2.6-3.4:1, apex acute to attenuate, base cuneate to obtuse and shortly decurrent down the petiole apex, slightly asymmetric, margin entire, lamina 0.28-0.62 mm thick, 0.76-1.22 mm at midrib, midrib conspicuous, raised on the adaxial and abaxial surfaces, grooved on adaxial; lateral nerves 16-25 on each side of the midrib, departing from midrib at 55-70 degrees, brochidodromous, looping 2-7 mm from margin, tertiary nerves conspicuous, raised abaxially, reticulate creating areolae; oil glands bright orange (transmitted light); in reflected light seen as “tiny purple spots” on adaxial surface when live, (*Wieringa* 248) or black spots (when dried), abaxially concolorous with blade, slightly raised on both abaxial and adaxial surfaces (dried), densest near margin, elliptic or oblong (-subcircular) 0.11-0.67 × 0.11-0.25 mm, 18-20 per 2 × 2 mm; 6-37 glands per areole; abaxial scales conspicuous (hand lense), bow-tie-shaped or subcircular, shortly 2-6 armed, red or reddish brown, 29-37 per 2 × 2 mm, 0.03-0.07 mm wide, adaxial surface glabrous; hairs absent on both surfaces. *Leaves* of plagiotropic stems as orthotropic, except with a smaller size range (leaves of orthotropic stems are 20-30% larger), petiole 6-11 × 2.6-3.4 mm, lamina 8.3-22.5 × 3.9-7.3 cm. *Inflorescences* axillary, present only on the distal half of the plagiotropic stems, in 3-5 successive nodes, each 3-14-flowered; peduncle 1-4 mm long; rachides 2-4 mm long; bracts caducous, ovate-lanceolate, 1.4-2.5 × 0.5-1.5 mm. Pedicel bases exserted from the rachis, in life “pink-red swollen towards the fruits” (*Wieringa* 248), when dried straw-coloured, floccose on rachides and bracts; pedicels descending, terete, thicker distally, 8-10 × 0.6-0.8 mm, glabrescent, oil glands at surface, red-brown to black sparse, raised, subcircular, elliptic or oblong, 0.11-0.45 × 0.09-0.18 mm.

Flower buds ovoid, 4.5-5.5 × 2.25-2.5 mm, apex acute; calyx cupular, lobes 5, overlapping to the right, ovate, 1.5-1.8 × 0.1-1.6 mm, apex acute to obtuse, oil glands covering c. 30% of calyx, orange to dark red-brown (reflected light), raised on outer surface, subcircular to oblong, 0.05-0.18 x 0.05-0.12 mm, margin ciliate, ciliae translucent, c. 0.09 mm long; corolla lobes overlapping and twisting to the right. Flowers 5-merous, pendulous, calyx as bud except lobes spreading to reflexed; Corolla spreading, pinkish in life (*F*.*J. Breteler* 12910), c. 9 mm diam.; tube shortly cylindric, adnate to staminal tube; corolla lobes ovate, c. 5 x c. 2.5 mm, apex acute, oil glands c. 15% of surface, visible and raised on both sides, red-brown (reflected light), subcircular, elliptic or oblong, 0.06-0.69 x 0.06-0.18 mm. Androecium erect, staminal tube shortly cylindric, c. 0.7 x c. 1.8 mm, filaments dorsiventrally flattened, c. 1.1 x c. 1 mm, widest at base; anthers narrowly elliptic, c. 3 x c.1.25 mm, apex acute, base sagittate, subbasifixed; oil glands forming an inconspicuous vertical line on the dorsal surface of both anther cells, and a clearer, sparser longitudinal line in the centre of the ventral surface along the connective, glands red brown to dark brown, subcircular, elliptic or oblong, 0.05 - 0.13 mm x 0.05 0.13 mm. Ovary subglobose, c. 1.1 x 0.9 mm, oil glands inconspicuous or absent; style terete, c. 4 mm long, oil glands covering c. 30% of surface, orange, subcircular to elliptic, c. 0.07 mm x c. 0.07 mm, stigma c. 0.1 mm. *Fruits* pendulous, dark red-brown, yellow (*J*.*J. Wieringa* 248), subglobose, 6-7 mm diam., calyx lobes spreading to reflexed, otherwise as flower, style base persistent, c. 0.4 x c. 0.2 mm; oil glands covering c. 40% of fruit surface, black, subcircular, elliptic or oblong, 0.15-0.9 x 0.15-0.27 mm, longest near calyx, glabrous; fruit wall c. 0.05 mm thick. Fig. 1 & 2.

**Fig. 2.**
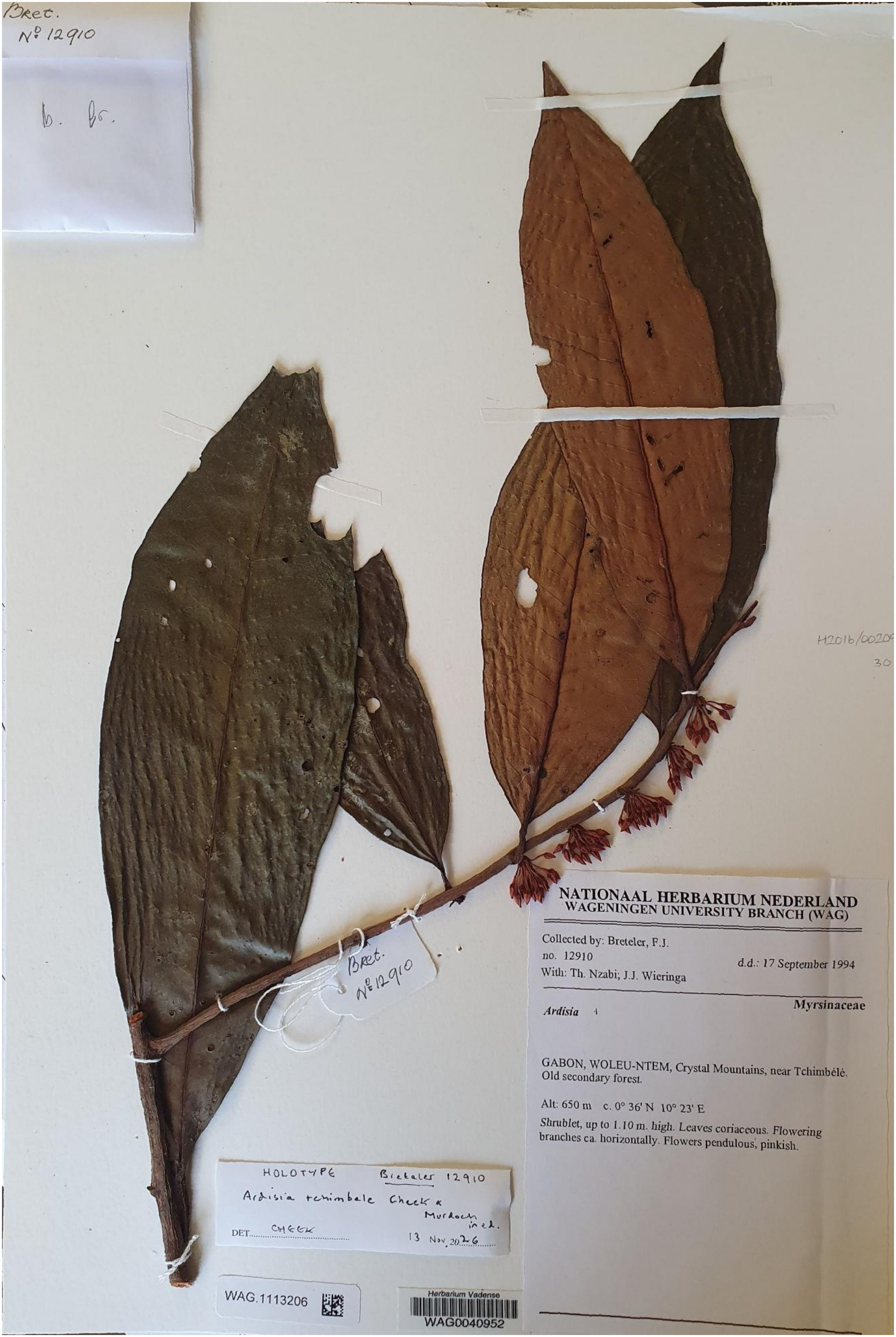
*Ardisia tchimbele*The holotype specimen (*Breteler* 12910, WAG).

### RECOGNITION

*Ardisia tchimbele* sp. nov. is similar to and has been identified as *A. midlbraedii* (Gilg & Schellenb.) de Wit, differing in having denser oil glands (500 per cm2 vs 60-200 cm2) which are raised on both adaxial and abaxial leaf surfaces (vs not raised), petiole 9-22 mm long (vs. 5-7 mm), longest leaf blade width 7.4-9.2 cm (vs. 5 cm), 16-25 lateral nerves (vs. 10).

### DISTRIBUTION

Gabon, Woleu-Ntem, Crystal Mountains, near Tchimbélé.**SPECIMENS EXAMINED. GABON**. Woleu-Ntem, Crystal Mountains, near Tchimbélé. Old secondary forest, 0°36’N, 10°23’E, 650 m, b., fr., 17 Sep. 1994, *F*.*J. Breteler* 12910 (BR, K, MO, WAG all iso., WAG holo.); Tchimbélé, 0.5 km. SE of Dam. Primary Forest. Steep hillside, 0°37’N, 10°24’E, fr., 18 Dec. 1989, *J*.*J. Wieringa* 248 (WAG !); 1 km. WNW of Tchimbélé. On bank of Bingiligwen river. Primary forest, 0°37’N, 10°23’E, 450 m, fr., 21 Apr. 1990, *J*.*J. Wieringa* 729 (WAG 1113202!; K!).

### HABITAT

Lowland evergreen rainforest; 460-650 m elev.

### CONSERVATION

The sites for this species are all downstream of the Tchimbélé hydroelectric dam constructed to supply the needs of Libreville, the capital c. 100 km distant. The reservoir behind the dam floods an area of former forest that extends c. 30 km upstream, and at its widest is c. 6 km when measuring flooded tributaries each side of the main stream. Since construction of the original dam in 1979, two further dams were opened or are under construction downstream (Anon 2025), flooding further areas of former forest to provide for the growing needs of the city, which also depends on the Crystal Mts for its water supply. Energy extraction from hydroelectric projects is most efficient and productive when the whole length of a stream is harnessed which can pose major threats for plants (Cheek et al. 2018), and can even result in the extinction of species, as was the case with *Saxicolella deniseae* Cheek (Cheek et al. 2022). Therefore, there is a risk that additional dam construction might create reservoirs that submerge one or all the known sites for *Ardisia tchimbele*. The georeferences for each of the three known specimen-sites (taken from the specimen labels) are placed at three of four corners of a square one minute (c. 1.8 km) apart, equating to an IUCN (2012) area of occupation of 4 km^2^ (using the IUCN required cell size). The extent of occurrence is estimated as being about the same size, and as all three sites are close to each other and have the same threat, we take them as a single threat-based location following the guidance of IUCN (2024). Therefore, we here provisionally assess the extinction risk of this species as Critically Endangered B1 + B2ab(i,ii,iii,iv,v). Should *Ardisia* sp. 1 be found to be conspecific with *A. tchimbele* (see Notes below), this would constitute a second location for the species and so the threat assessment would likely be lowered to that of Endangered.

### ETYMOLOGY

*Ardisia tchimbele* is named (noun in apposition) for the collection location of all the known specimens: in the area of the Tchimbélé dam near the village of that name.

#### NOTES

The isotype at BR has metadata on GBIF that incorrectly states “ [Ardisia_zenkeri](https://www.gbif.org/occurrence/1840275651) Gilg collected in Gabon by Meise Botanic Garden (licensed under CC BY 4.0)” (Meise Botanic Garden 2026). It does not state who has made the identification, while the actual specimen image featured is identified only to genus. Similarly with the MO isotype, GBIF states “Breteler - 12910 - Gabon - [Ardisia_Sw](https://www.gbif.org/occurrence/2268818954). collected in Gabon by MBG (licensed under http://creativecommons.org/licenses/by-nc-sa/3.0/)” (Teisher & Stimmel 2026). Also on GBIF the metadata for the WAG duplicate states “[Ardisia Sw](https://www.gbif.org/occurrence/2515820159). collected in Gabon by Naturalis Biodiversity Center (licensed under [CC0_1.0](http://creativecommons.org/publicdomain/zero/1.0/))” (Bijmoer et al. 2026). For the K duplicate GBIF states “[Ardisia_Sw](https://www.gbif.org/occurrence/5839108970). collected in Gabon by Royal Botanic Gardens, Kew (licensed under [CC_BY_4.0](http://creativecommons.org/licenses/by/4.0/))” (Royal Botanic Gardens, Kew 2026). In fact, Frans Breteler, the collector of this specimen, was working for Herbarium Vadense at Wageningen (WAG) at the time of collection and not any of these other institutions.

*Wieringa* 729 is a mixed collection of two or more very different species of *Ardisia*. Apart from *A. tchimbele* (the K and WAG duplicates), the duplicate at MO (barcode 6635202) is a different, monocaul taxon with oblanceolate leaves, in fruit, the GBIF image being insufficient to identify it to species using Murdoch & Cheek (2026a) which revises this group in Gabon. The BR duplicate (barcode 0000017904896) may be the same taxon as that of MO, or a third taxon. *Wieringa* 729 was not previously identified to species and was not cited in Sosef *et al*. (2006), while *F*.*J. Breteler* 12910 was cited in Sosef *et al*. (2006) as an *Ardisia* but without a species determination. *Wieringa* 248 was cited in Sosef *et al*. (2006) as *Ardisia mildbraedii*, det. Wieringa 1990.

*Ardisia tchimbele* keys out (see Introduction) as, and bears resemblance to Taton’s *Ardisia* sp.1 (Taton 1979), for which he cites only one specimen (*N. Hallé & J. F. Villiers* 4739) also from Gabon’s Crystal Mountains, at Nkam, which is only 37 km distant from the location of the three accepted specimens. Taton’s brief description of *Ardisia* sp.1 is largely in line with our description of *A. tchimbele* above. We have inspected the image of the specimen on GBIF. It differs from *A. tchimbele* in having an undulate, not an entire, leaf margin.

Therefore, we hesitate to include it as this species. Further, we have not studied the leaf oil gland characteristics, nor the characteristics of the abaxial leaf scales. Should these prove to be identical with those of *A. tchimbele*, and should no additional differential characters be detected, we would likely include this specimen as *A. tchimbele* despite the difference in leaf margin, interpreting this as part of the natural variation within *A. tchimbele*.

## Discussion

### The Crystal Mountains

The Crystal (in French Cristal) Mountains of Gabon are relatively well known as a centre of plant diversity in Gabon (see Breteler 1990, De Wilde 1994), as a Pleistocene Refuge (Sosef 1994) and for having endemic plant species (Bissiengou & Sosef 2008). The range runs roughly parallel to the coast from which it is separated by a coastal plain of c. 50 km wide with an elevation of c. 50 m. The mountains form an often ill-defined mass c. 50 km wide w to E and extend c. 75 km north of the Ogooué River, crossing the border into Rio Muni in the north. They are of ancient basement rocks, with irregular mostly N:S ridges often reaching 700 m (rarely exceeding 800 m) elevation and valleys. 800 m elevation is considered the threshold at which submontane forest begins, at least in neighbouring Cameroon (Cheek et al. 2004; Murphy et al. 2023) and therefore the vegetation is considered as mainly lowland evergreen forest, with few areas of open habitat. Inland of the mountains, elevation drops but is still in the area of 500 m elevation or more.

Botanical exploration in Gabon began in the 18^th^ century (Middleton s.n. collected in 1787, type of *Cryptacanthus* I. Darbysh. & Kiel., Acanthaceae, Darbyshire et al. 2025). But botanical surveys of the Crystal Mts began many decades later, in 1861 by Gustav Mann, whose collections include the point endemic *Ardisia bracteata* Baker from the border of Gabon with Rio Muni which has not been recollected in more than 160 years (Murdoch & Cheek 2026a).

According to Bissiengou & Sosef (2008), at least 29 plant species are already known to be endemic to the Gabonese part of the Crystal Mountains among which are four species of *Begonia* (*B. aggeloptera* N.Hallé, *B.erectocaulis* Sosef, *B. erectotricha* Sosef, and *B. peperomioides* Hook.f., fide Sosef 1994), four orchids and seven Rubiaceae. To these they added the shrub *Campylospermum louisii* Bissiengou & Sosef (Ochnaceae). Amongst the additional endemic species described recently are *Uvariopsis niangadoumae* Couvreur & Dagallier (Couvreur et al. 2025), *Uvariopsis citrata* Couvreur & Niang. (Couvreur et al. 2016), and even an endemic genus *Sirdavidia solannona* Couvreur & Sauquet, (all Annonaceae, Couvreur et al. 2015). *Ardisia mela* Cheek & Murdoch (Murdoch & Cheek 2026b), endemic to Méla is in the course of formal publication. There is no doubt that many additional new species including point endemics remain to be found and published from the Crystal Mountains, e.g. two new unpublished species of *Afrothismia* (Afrothismiaceae, Cheek et al. 2024).

The Cristal Mountains are partly protected by the Parcs National des Monts de Cristal (Crystal Mts National Park, Anon continuously updated), which are divided into two separate blocks, Mt Sene and Mt Mbe, the total area of which is 1,200 km^2^ (Anon, continuously updated). However, very large areas, including that which hosts *Ardisia tchimbele*, remain unprotected.

## Conclusion

Publication of new species to science is important to improve the possibility of their conservation (Cheek et al. 2020) which has become urgent since three out of four species published in recent years are threatened with extinction at the point of publication (Brown et al. 2023). Although Gabon is blessed with the highest coverage of intact original forest of all countries in tropical Africa, and the level of threats seem relatively low, nevertheless there are cases of apparent species extinction of Gabonese endemic species, such as *Pseudohydrosme buettneri* Engl. and *P. bogneri* Cheek & Moxon-Holt (both Araceae, respectively Cheek et al. 2021 and Moxon-Holt & Cheek 2021). We advise that a survey of *Ardisia tchimbele* is carried out to collect more detailed information on threats, on the range, regeneration and number of individuals of the population, so as to inform a species conservation action plan (e.g. Couch et al. 2022). Options should include the area being considered in an Important Plant Area (Darbyshire et al. 2017) and also *ex situ* conservation.

## Acknowledgements

We acknowledge the efforts of the lead collectors of the specimens of this new species, Jan Wieringa and Frans Breteler. Without their work in making specimens and recording field data, this paper would not exist. We especially thank Corinne Sarthou-Gasc of the curator team at Herbier National, Muséum national d’histoire Naturelle, Paris for superb assistance accessing specimens of *Ardisia* at P in July 2025 for our research visit there, and also Germinal Rouhan. We also thank Mark Carine and the staff of BM, and the curator-botanist team at K, led by Renata Borosova. Gaia Caravella is thanked for the line drawing. HM thanks fellow student Caitlyn McDaniel for discussions and collaboration on *Ardisia* in 2025, during which this paper was completed as part of an MSc Project on the R.B.G., Kew-Queen Mary University of London Plant and Fungal Taxonomy, Diversity and Conservation.

## Data availability statement

This article has no additional data.

